# Ras inhibition by trametinib treatment in *Drosophila* attenuates gut pathology in females and extends lifespan in both sexes

**DOI:** 10.1101/356295

**Authors:** Jennifer C Regan, Yu-Xuan Lu, Ekin Bolukbasi, Mobina Khericha, Linda Partridge

## Abstract

Females of most species live longer than do males. Furthermore, lifespan-extending
interventions in laboratory model organisms are often more effective in females (Regan and Partridge 2013). For instance, genetic and pharmacological suppression of activity of the insulin/insulin-like signalling - target of rapamycin (IIS-TOR) network generally extends female lifespan more than that of males in both *Drosophila* and mice (Clancy et al. 2001; Selman et al. 2009). We previously showed that attenuation of Ras-dependent IIS signalling by treatment with the FDA-approved MEK inhibitor, trametinib extends lifespan in females (Slack et al. 2015). Here, we demonstrate that trametinib treatment has beneficial effects on female-specific, age-related gut pathologies, similar to those obtained through dietary restriction (Regan et al. 2016). Importantly, we identify Ras inhibition as an effective lifespan-extending manipulation in males as well as females, pointing to parallel mechanisms of lifespan extension by trametinib in both sexes.

## Introduction

Lifespan extending manipulations affecting the insulin/insulin-like signalling - target of rapamycin (IIS-TOR) network have variable relative efficacy in females and males. For example, broadly targeting the network through dietary restriction (DR) is less effective at extending median lifespan in *Drosophila* males than in females (Magwere, Chapman, and Partridge 2004; Regan et al. 2016), whereas heterozygotes for the null mutant of the insulin receptor substrate homolog, *chico*, show extended lifespan in both sexes (Clancy et al. 2001). There is considerable interest in understanding the mechanisms underpinning sexually dimorphic responses to lifespan-extending manipulations, in order to understand their mechanisms and to target them appropriately in the general population.

We recently showed that the FDA-approved MEK inhibitor trametinib significantly extends lifespan in female *Drosophila melanogaster* (Slack et al. 2015). Ras inhibition contributes to lifespan extension from reduced insulin/IGF-1 (IIS) signalling, and pharmacological inhibition of the Ras pathway using trametinib is effective in extending lifespan without several of the negative effects inherent in IIS-inhibition (Slack et al. 2015). However, mechanisms for extension of life by trametinib remain elusive.

The effect of trametinib on male lifespan has not been described. Here, we show that male lifespan is significantly extended by trametinib, but this effect is considerably more variable in males than in females. Treatment in females results in a strong attenuation of intestinal pathology revealed by analysis of ageing gut phenotypes. Age-related intestinal pathology does not usually develop, or develops at a much slower rate, in control males (Regan et al. 2016) and is thus unaffected by drug treatment. Our study identifies Ras inhibition as an effective lifespan extending manipulation in males as well as females, and importantly points to parallel mechanisms of lifespan extension by trametinib in both sexes, in addition to its beneficial effects in intestinal tissue in females.

## Trametinib extends lifespan in both sexes, but more robustly in females than males

We treated adult flies of both sexes, throughout their lifespan, with titrated concentrations of trametinib added to the food medium. The dose that previously gave a maximum extension in females, 15.6μM (Slack et al. 2015), was designated ‘1x’. Male flies have been shown to eat less than mated female flies by various methods (Deshpande et al. 2014; Wong et al. 2009), but may be differently sensitive to drug treatment, therefore flies of both sexes were also fed a 0.1x (1.56μM) and 2x (31.2μM) dose. As we have previously shown (Slack et al. 2015), female lifespan was extended significantly at all doses (0.1x = 5.1% extension, *p*=0.0004; 1x = 14.7% extension, *p*=3.69×10^−24^; 2x = 14.7% extension, *p*=3.86×10^−21^), although with some early mortality at the highest (2x) dose (Fig 1A). Male median lifespan was extended at all doses, although extension at the lowest dose (0.1x) was not significant. Male lifespan was extended to a slightly lesser extent than female (0.1x = 6.1% extension, *p*=0.43; 1x = 12.8% extension, *p*=0.0013; 2x = 12.8% extension, *p*=1.27×10^−6^ (Fig 1). To compare the extent of lifespan extension in the two sexes, we used Cox Proportional Hazards (CPH) analysis, where the interaction between trametinib treatment and gender was found to be significant (*p*=0.0011), indicating that the sexes respond to treatment with trametinib to a different degree (Table S1). Male lifespan extension was also less reliably extended than that of females, with no extension at any dose in 1 out of 3 trials (Fig 1 and Table S1).

**Figure 1.**
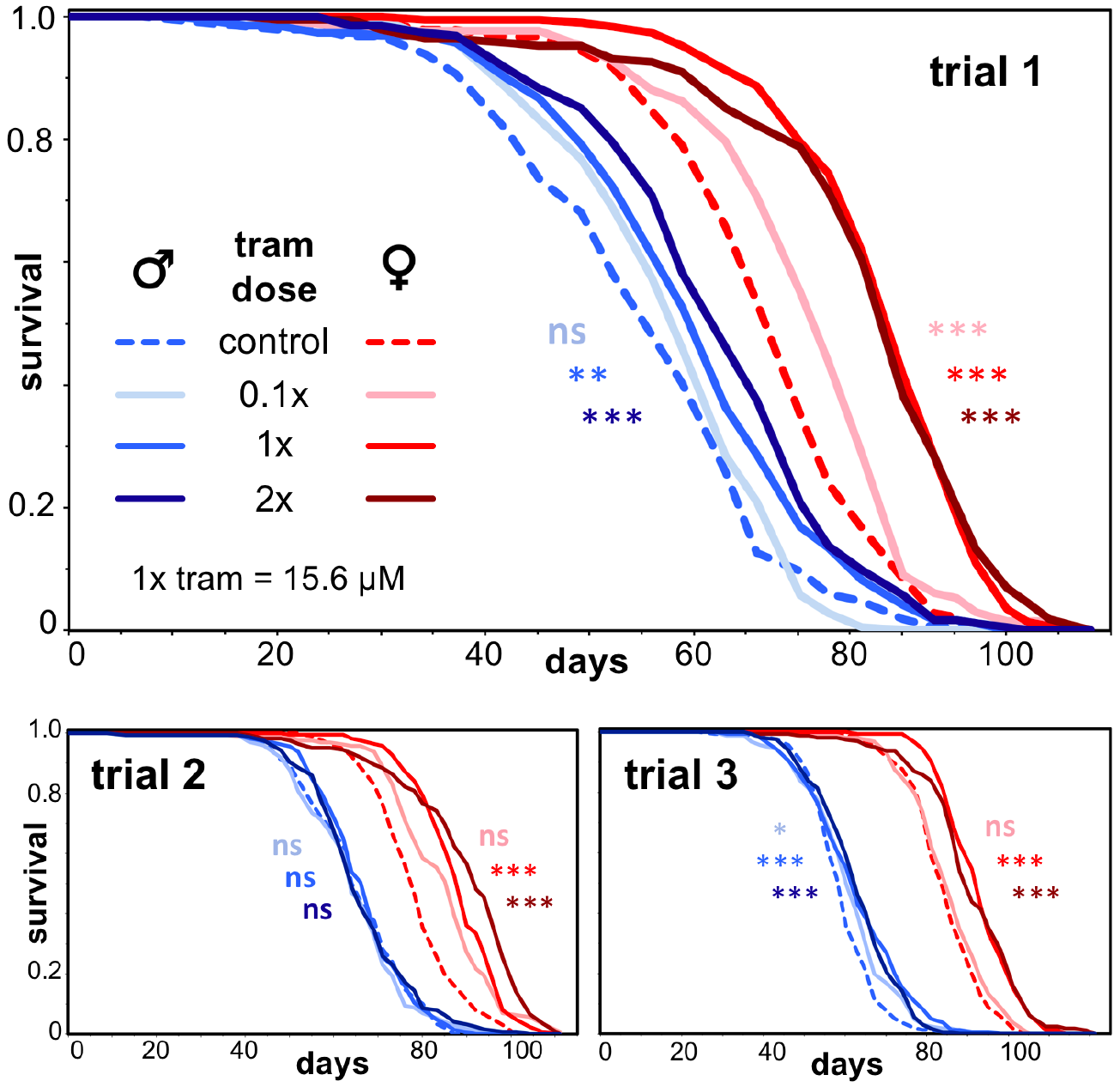
Chronic treatment with trametinib extends lifespan in males and females. (Females: 0.1x = 5.1% extension, *p*=0.0004; 1x = 14.7% extension, *p*=3.69×10^−24^; 2x = 14.7% extension, *p*=3.86×10^−21^. Males: 0.1x = 6.1% extension, *p*=0.43; 1x = 12.8% extension, *p*=0.0013; 2x = 12.8% extension, *p*=1.27×10^−6^). Differences in survival were tested by Log-rank; *p* values given are for trial 1. For CPH analysis of Trial 1 and a summary of all trials and *n*, see Supplementary Table 1. Tram = trametinib. 0.1x = 1.56 μM, 1x = 15.6 μM, 2x = 31.2 μM. * *p* ≤ 0.05; ** *p* ≤ 0.01; *** *p* ≤ 0.001.

## Trametinib represses intestinal stem cell (ISC) division in both sexes

Trametinib is an FDA-approved drug for the treatment of melanoma. It is a small molecule inhibitor of Mek kinase, preventing the Ras-dependent activation of Erk. Ras regulates cell proliferation and survival, and over-activation of Ras is highly oncogenic, with Ras mutations found in over a third of human tumours (Stephen et al. 2014). In addition, EGFR/Ras/Erk/ETS signalling is known to be involved in the regulation of ISC division and tumour formation in adult flies (Biteau and Jasper 2011; Jin et al. 2015; Patel, Dutta, and Edgar 2015; Xu et al. 2011), and trametinib reduced mortality in a *Drosophila* Ras-Pten cancer model (Levine and Cagan 2016). Therefore, we were surprised to find that trametinib treatment did not influence age-related increases in ISC division (Slack et al. 2015). When we analysed ISC division in trametinib-treated flies of both sexes in greater numbers and at earlier time points, we found that ISC activity was suppressed in guts from trametinib-treated flies (Fig 2A). This represented a strong attenuation in female guts, which have a higher ISC division rate than males (Hudry, Khadayate, and Miguel-Aliaga 2016; Jiang et al. 2009; Regan et al. 2016). A significant suppression was also observed in males, despite the low basal rate of division (Fig 2A).

**Figure 2.**
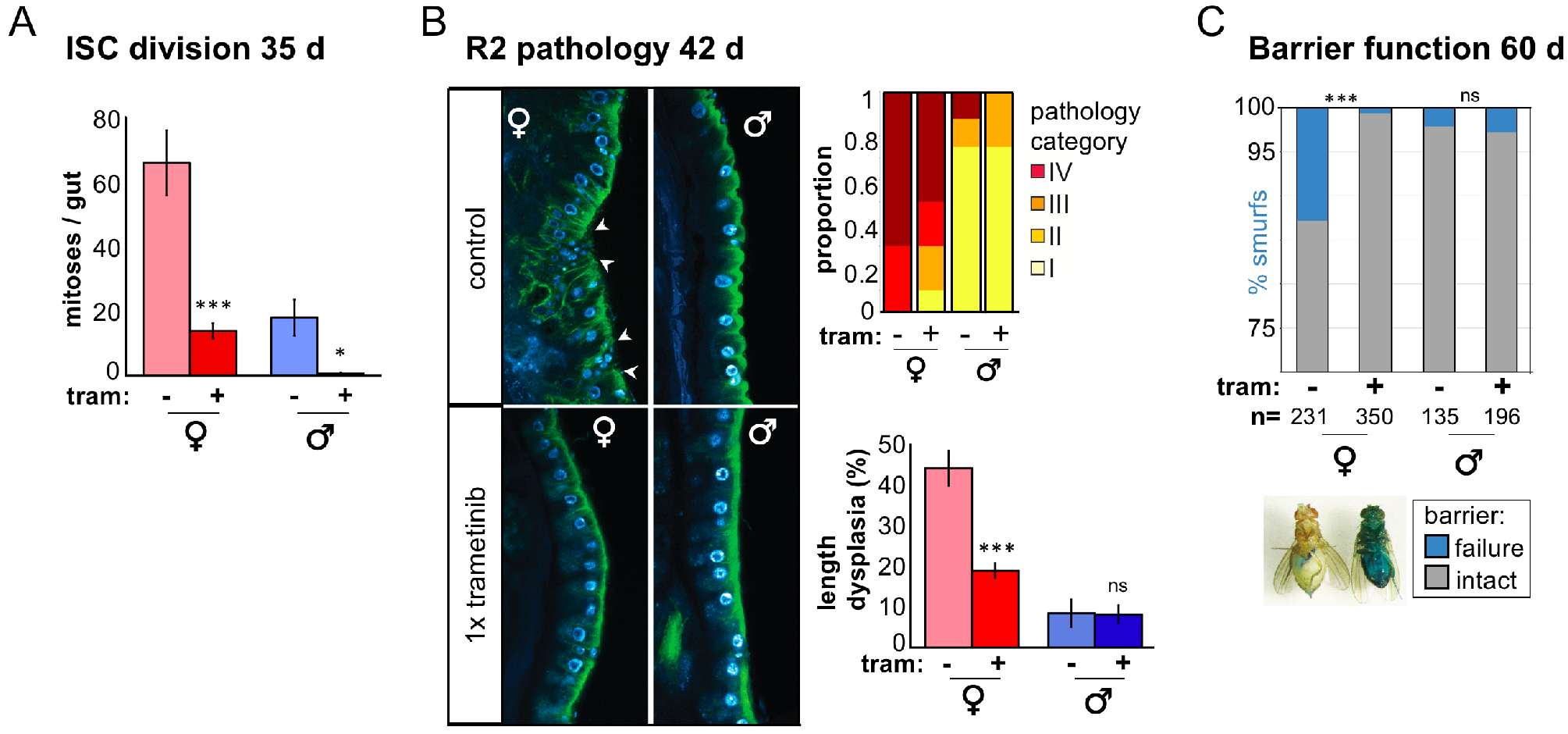
(A) Treatment with trametinib reduces intestinal stem cell division in 35 d males and females. Mitoses were measured by positive staining with the phosphohistone H3 antibody, and scored in whole midguts. Data are representative of 2 independent experiments. Sample sizes for each condition as follows: female DMSO, n=29; female trametinib, n=26; male DMSO, n=10; male trametinib, n=10. Tram = trametinib. 1x = 15.6 μM. * *p* ≤; 0.05; ** *p* ≤ 0.01; *** *p* ≤ 0.001. (B) Treatment with trametinib reduces intestinal epithelial pathology in females. Intestines from *Resille-GFP* males and females were imaged, and pathology scored and binned into categories. (I = no pathology; II = sporadic small tumours; III = epithelial wounds and small tumours; IV = large tumours, severely disrupted epithelium). Percentage length dysplasia was blind-scored from luminal sections of approx. 800μm of the R2 region of trametinib and control treated intestines. Sample sizes for each condition as follows: female DMSO, n=10; female trametinib, n=9; male DMSO, n=8; male trametinib, n=8. Treatment with trametinib ameliorates intestinal barrier function decline in 60 d females (*p* ≤ 0.0001). Males were not significantly affected (*p* = 1.0). Data is representative of 2 independent experiments. Sample sizes for each condition as follows: female DMSO, n=231; female trametinib, n=350; male DMSO, n=135; male trametinib, n=196.

## Trametinib attenuates age-related intestinal pathology and loss of gut barrier function in females

It is not clear to what extent ISC division *per se* regulates lifespan; although it is certainly strongly linked with longevity (Biteau et al. 2010; Rera et al. 2011), recent studies have shown that homeostasis of differentiated cells is important for maintenance of gut function, including barrier function (Bolukbasi et al. 2017; Resnik-Docampo et al. 2016). Therefore, we analysed pathology of the epithelium, and barrier function of the intestine in 1x trametinib-treated flies over age. Using high-resolution imaging of the R2 region of guts carrying a fluorescent epithelial marker (*Resille-GFP*), we found that female flies treated with trametinib were less likely to develop the strongest pathologies (categories III and IV, i.e. intestines carrying large tumours and with significant epithelial disruption; Fig 2B). To perform more quantitative analysis, we measured proportion of dysplasia in standardized luminal sections in the same intestines. Trametinib-treated females had a significantly lower proportion of their intestines developing a dysplastic phenotype (Fig 2B). Males were not affected by trametinib treatment (Fig 2B), probably because levels of age-related dysplasia are low in untreated, control males ((Regan et al. 2016) and this study).

When we measured barrier function by feeding the flies a non-absorbable blue dye (Rera, Clark, and Walker 2012), we found that loss of barrier integrity was strongly suppressed in females treated with 1x trametinib (Fig 2C). Again, this is in contrast with our previous results (Slack et al. 2015), and may reflect insufficient statistical power. No significant difference was found comparing treated and untreated males, which, in the *w*^*Dah*^ outbred line, have a very low rate of barrier function loss at 60 d (<5%; (Regan et al. 2016) and this study; Fig 2C), despite this time point being closer to the median lifespan of males than of females (Fig 1).

## Discussion

Here, we show that treatment with the Mek-inhibitor trametinib significantly extended lifespan in both male and female *Drosophila*. However, the magnitude of lifespan extension was greater in females than in in males. In addition, we found that it was less reliable in males, where male lifespan was not extended at all in one of three trials. We do not know the source of this variability, but we can speculate that it was due to interaction of the drug with an environmental factor such as dietary components or infection. In the case of enteric infection, inhibition of ISC division by trametinib may cross a critical minimal threshold for males where they are unable to repair their intestines, suppressing lifespan extension by the drug.

Given that trametinib is an FDA-approved drug, and we have shown it can extend lifespan in both sexes in our *Drosophila* model, there is considerable motivation to understand its mechanism for promoting longevity. Here, we show that trametinib treatment significantly ameliorated age-related gut pathologies in females, such as tumour formation and loss of barrier function, which are known to be limiting to female lifespan. We suggest that this may be important for the lifespan extension in females afforded by trametinib treatment, and may account for the more robust efficacy in females compared to males.

We hypothesise that trametinib has other significant, parallel functions that promote longevity besides its effects on gut pathology. Males do not develop significant gut pathologies, at least those we can measure through observation of the epithelium, analysis of tumour formation and barrier function against ingested dye. Our previous work suggests that manipulations that act mainly to reduce gut decline are strongly sexually dimorphic in their effect on lifespan (Regan et al. 2016), but this does not appear to be the case with trametinib.

Trametinib may function through the regulation of intestinal cell function, such as ISC activity or signalling, although the quiescent nature of ISCs in males somewhat argues against this possibility. The transcription factor *Aop* is both necessary and sufficient to mediate the beneficial effects of attenuated Ras signaling (Slack et al. 2015), and overexpression of *Aop* in the fat body is sufficient to extend lifespan (Alic et al. 2014). These results are suggestive of a role for trametinib in fat body functions that are important for longevity in both males and females.

In the drive to develop personalised, anti-ageing therapeutics, understanding the influence of sex is key. Investigating sex-specific responses to interventions both reveals relative efficacies, and gives insight in to underlying mechanisms of action.

## Experimental Procedures

### Fly husbandry and strains

Stocks were maintained and experiments conducted at 25°C on a 12-hr light/dark cycle at 60% humidity, on food containing 10% (w/v) brewer’s yeast, 5% (w/v) sucrose, and 1.5% (w/v) agar. Food was supplemented with Trametinib (LC Laboratories) from a 62.4 mM stock solution in DMSO at the following concentrations; 1.56 μM (0.1 x), 15.6 μM (1 x), 31.2 μM (2 x); maintaining a final DMSO concentration of 0.05% (v/v). For control treatments, equivalent volumes of the vehicle alone were added. For measurement of gut histology and lifespans, flies were reared at standard larval density and eclosing adults were collected over a 12-hr period. Flies were mated for 48 hr before being sorted into experimental vials at a density of 10 flies per vial. Flies were transferred to fresh vials thrice weekly, and for lifespan, deaths/censors were scored during transferral. All experiments were performed on the ‘outbred’ (cage-maintained) *w*^*Dahomey*^ (*w*^*Dah*^) line except for gut pathology analysis, which used (*Resille-GFP;* FBti0141278) backcrossed for six generations into *w*^*Dah*^. *Resille-GFP* was originally obtained from the Flytrap project (Kelso et al. 2004) and was a gift from A. Jacinto.

### Imaging of gut pathology

Guts were dissected from live flies in ice-cold PBS and immediately fixed in 4% formaldehyde for 15 min. Guts were mounted in mounting medium containing DAPI (Vectastain) and endogenous GFP was imaged immediately. Images were captured with a Zeiss (UK) LSM 700 confocal laser scanning microscope using a 40x oil- immersion objective.

### Immunohistochemistry

The following antibodies were used for cell division analyses; primary antibodies: rabbit anti-PH3 (Cell Signaling (Danvers, MA) 9701) 1:500; mouse anti-GFP (Cell Signaling 2955) 1:1000. Secondary antibodies: Alexa Fluor 594 donkey anti-rabbit ((A21207) Thermo Fisher Scientific, Waltham, MA) 1:1000; Alexa Fluor 488 donkey anti-mouse (A21202) 1:1000. Guts were dissected in ice cold PBS and immediately fixed in 4% formaldehyde for 15 min, serially dehydrated in MeOH, stored at - 20 °C, and subsequently stained. Guts were washed in 0.2% Triton-X/PBS, blocked in 5% bovine serum albumin/PBS, incubated in primary antibody overnight at 4 °C and in secondary for 2 hr at RT.

### Gut barrier analysis (Smurf assay)

Gut barrier efficiency was analyzed by placing flies on blue food prepared using using 2.5% (w/v) FD&C blue dye no. 1 (Fastcolors) as previously described (Rera et al., 2012), except flies were kept on blue food for 48 hr before phenotype was scored. N.B. This experiment was performed twice, in parallel with lifespan trials 1 and 3 (Fig 1), both of which extended lifespan in both sexes.

### Statistical analysis

Statistical analysis was performed in Excel (Microsoft), Prism (Graphpad) or R. Survival data were analysed with either Log-rank test using Excel (Microsoft) or Cox Proportional Hazards with R using the survival package (Terry Therneau, https://cran.r-project.org/web/packages/survival/index.html). Gut pathology data were analysed using Student’s *t* test or Fisher’s Exact test. Sample sizes are indicated in figure legends.

## Acknowledgements

We thank D. Menden, I. Snoeren, M. Ahmad and R. Herzog for technical assistance. We thank N. Alic, A. Dobson and C. Slack for useful discussion and comments on the manuscript. We acknowledge funding from a Wellcome Trust Strategic Award (WT081394) and the Max Planck Society. The research leading to these results has received funding from the European Research Council under the European Union’s Seventh Framework Programme (FP7/2007-2013)/ERC grant agreement number 268739 to L.P and J.R.

